## Supplementary figures and images for "Tmem263 deletion disrupts the GH/IGF-1 axis and causes dwarfism and impairs skeletal acquisition"

### Figure 2 -figure supplement 1

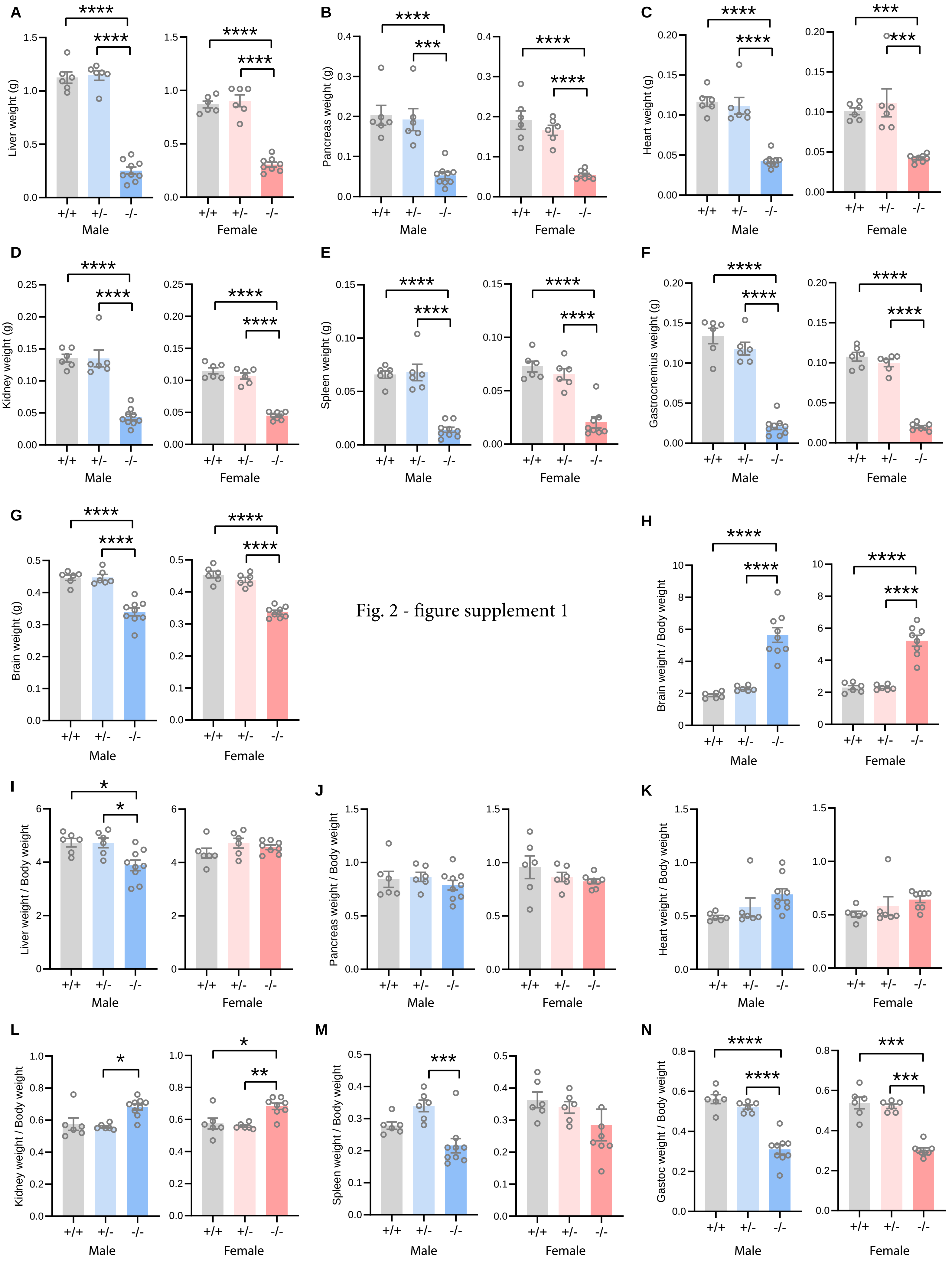


Fig. 2 - figure supplement 1

### Figure 6-figure supplement 1

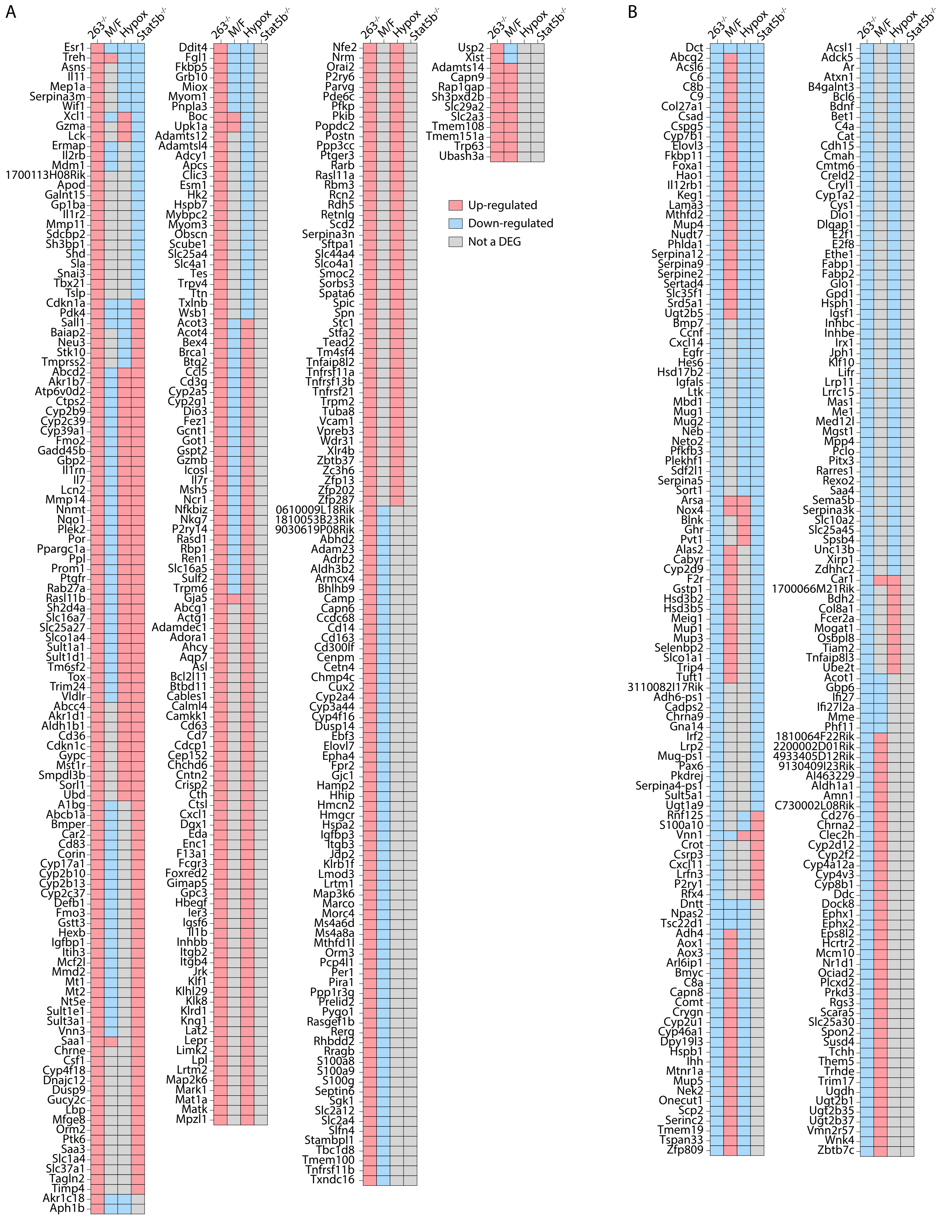
